# *Smed-egfr-4* is required for planarian eye regeneration

**DOI:** 10.1101/446013

**Authors:** Elena Emili, Maclà Esteve Pallarès, Rafael Romero, Francesc Cebrlà

## Abstract

Planarians are amazing animals that can regenerate a whole body from a tiny piece of them thanks to their pluripotent stem cells, the neoblasts. Planarian neoblasts include both pluripotent stem cells and specialized lineage-committed progenitors that give rise to all the mature cell types during regeneration and homeostatic cell turnover in these plastic animals. Little is known, however, about the mechanisms that regulate neoblast differentiation. Recently, it has been shown that *Smed-egfr-1,* a homologue of the epidermal growth factor receptor (EGFR) family is required for the final differentiation of the gut progenitors into mature cells but not for their specification. As planarians have several EGFR homologues it has been proposed that they could have diverged functionally to regulate the differentiation of the different cell types found in these animals. Here, we report on the function of *Smed-egfr-4* on eye regeneration. The silencing of this gene by RNAi results in animals regenerating smaller eyes compared to controls. The numbers of both eye mature cell types, photoreceptor neurons and eye-cup pigment cells, are significantly decreased in the *Smed-egfr-4(RNAi)* animals. In contrast, the number of eye progenitor cells expressing the specific markers *Smed-ovo* and *Smed-sp6-9* is increased. These results suggest that *Smed-egfr-4* would be required not for the specification of eye progenitor cells but rather for their final differentiation and support the idea that in planarians the EGFR pathway could play a general role regulating the differentiation of lineage-committed progenitors.

## Introduction

Unravelling the mechanisms that regulate stem cell differentiation is pivotal to understand the amazing regenerative capabilities shown by some animals that maintain stem cells throughout their lives and that can be activated after an injury or amputation. Freshwater planarians are one of such classical model for stem cell-based regeneration (for recent reviews, see Cebrià et al., 2018; Rink 2018). Planarian neoblasts are an heterogeneous cell population that includes truly pluripotent undifferentiated neoblasts, referred as cNeoblasts, (Wagner et al., 2011; Zeng et al., 2018), as well as specialized lineage-committed progenitors specific for the different mature cell types (Scimone et al., 2014; Zhu and Pearson, 2016). Although these cell-specific progenitors have been largely characterized for most cell types relatively little is known about the genes and signalling pathways that regulate both the specification of these progenitors from the pluripotent stem cells and their subsequent final differentiation into mature cells (Zhu et al., 2015; Solana et al., 2013; Tu et al., 2015). Recently, it has been suggested that the epidermal growth factor receptor (EGFR) signalling pathway could have an important general role during neoblast differentiation (Barberán and Cebrià, 2018). Thus, the silencing of *Smed-egfr-1* results in defects in the regeneration and maintenance of the gut (Barberán et al., 2016a). After *Smed-egfr-1* RNAi there is a decrease in the number of newly differentiated mature gut cells accompanied by an increase in the number of gut progenitors that fail to differentiate. Therefore, *Smed-egfr-1* appears to be necessary not for the specification of the gut progenitors but for their final differentiation (Barberán et al., 2016a). In planarians, the EGFR family has been expanded and 6 homologues have been identified in the model species *Schmidtea mediterranea* (Barberán et al., 2016b). These different EGFRs show distinct expression patterns and have been suggested to be involved in the differentiation of different cell types such as neurons or protonephridia (Fraguas et al., 2011; Rink et al., 2011). Based on the function of *Smed-egfr-1* in gut differentiation, the fact that the different EGFRs are expressed in different tissues and cell types and that some EGFRs have been implicated in cell differentiation, a current model proposes that EGFRs might have a general function in the final differentiation of different populations of lineage-committed progenitors (Barberán and Cebrià, 2018). Here, we report on the function of *Smed-egfr-4* on eye regeneration. The silencing of *Smed-egfr-4* results in smaller eyes formed by a reduced number of photoreceptor neurons and pigmented eye-cup cells. Interestingly, this reduction in the number of mature eye cells correlates with an increase in the number of eye progenitors. Therefore, these results suggest that *Smed-egfr-4* is required for the final differentiation of eye-progenitor cells and would support a general role of EGFR genes in neoblast differentiation in planarians.

## Results

### Smed-egfr-4 is expressed rather ubiquitously including the eyes

Whole-mount *in situ* hybridizations showed that *Smed-egfr-4* was expressed in the cephalic ganglia, pharynx and in the mesenchyme around the gut branches (Fig. 1A). Also, a faint expression was detected in the photoreceptor cells (Fig. 1B). The expression in the mesenchyme did not appear to correspond to neoblasts as the expression pattern of *Smed-egfr-4* was not reduced in irradiated animals in which the neoblast population had been eliminated (Fig. 1C). In order to analyse the expression of *Smed-egfr-4* during regeneration planarians were decapitated and the regeneration of a new head was monitored. *Smed-egfr-4* was expressed in the regenerative blastema from day 1 after amputation (Fig. 1D). From day 2-3 *Smed-egfr-4* was also detected in the newly formed brain primordia (Fig. 1 E-F) and its expression was maintained high within the blastema and in the regenerating cephalic ganglia throughout the whole process of regeneration (Fig. 1 G-I).

**Fig. 1.**
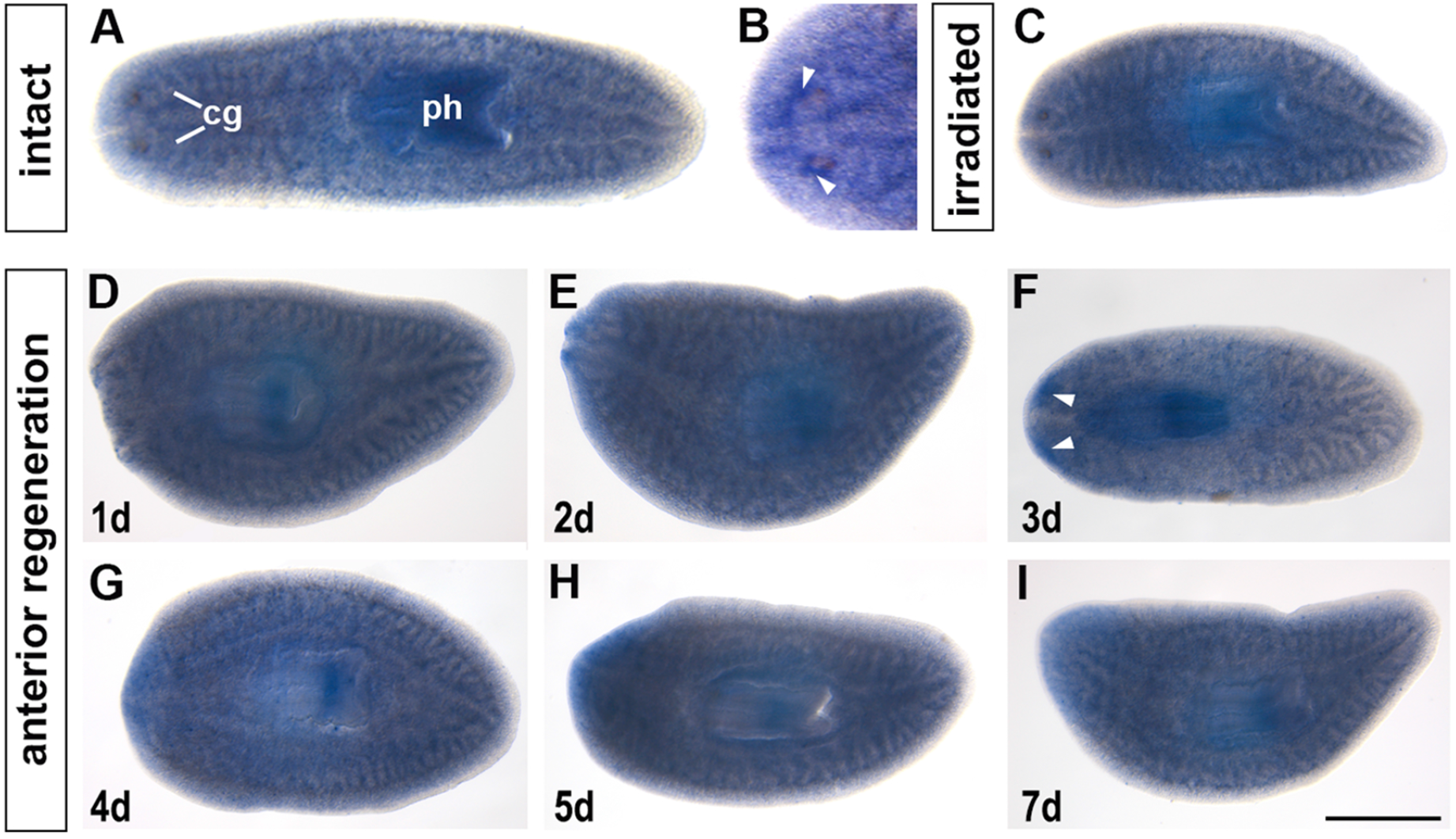
*Smed-egfr-4* expression pattern in intact and regenerating planarians. **(A,B)** *Smed-egfr-4* is expressed in the CNS, pharynx, mesenchyme and in the photoreceptors (arrowheads in B). (C) Irradiation does not alter the expression pattern of *Smed-egfr-4.* (D-I) *Smed-egfr-4* expression during anterior regeneration. Arrowheads in (F) point to the new brain primordia. cg, cephalic ganglia, ph, pharynx. Scale bar, 400 μm.

### Smed-egfr-4 silencing does not affect CNS regeneration

In order to analyse the function of *Smed-egfr-4* we performed RNAi-based functional experiments. As *Smed-egfr-4* was expressed in the central nervous system (CNS) we first analysed whether its silencing resulted in defects in the regeneration of the cephalic ganglia. Whole-mount *in situ* hybridizations for different neural specific markers revealed that *Smed-egfr-4* RNAi animals could regenerate a proper CNS with normal patterns for the distinct neuronal subpopulations analysed (Fig. 2). Those included GABAergic neurons *(Smed-gad),* dopaminergic neurons *(Smed-th)* and the brain lateral branches *(Smed-gpas).*

**Fig. 2.**
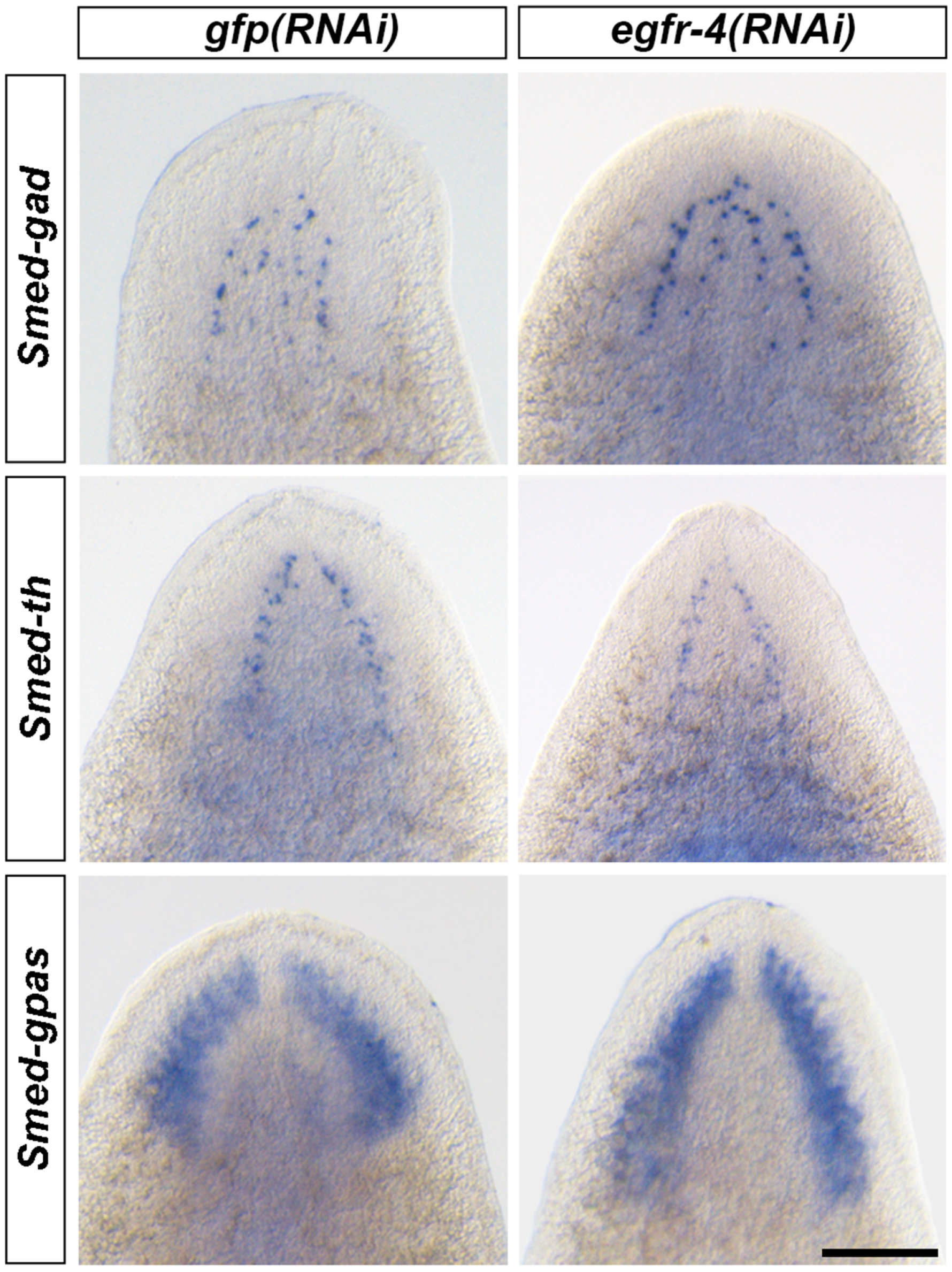
Normal CNS regeneration after *Smed-egfr-4 RNAi.* After 11 days of regeneration, *Smed-egfr-4(RNAi)* animals regenerate normal patterns of different neuronal populations. Scale bar, 150 μm.

### The silencing of Smed-egfr-4 results in the regeneration of smaller eyes

Two cell types constitute planarian eyes: photoreceptor neurons and pigmented eye-cup cells that can be easily distinguished in live animals. The photoreceptor neurons are located in the whitish area adjacent to the pigmented eye-cup (Fig. 3). *Smed-egfr-4(RNAi)* planarians regenerated smaller eyes compared to controls. In order to have a quantitative support for these apparent differences, and taking into account that there is a correlation between the size of the head and the size of the eyes (bigger heads have bigger eyes and smaller heads have smaller eyes), we measured the ratio between the eye length and the head length from the posterior base of the eyes to the most anterior tip of the head. Those measurements supported that the silencing of *Smed-egfr-4* resulted in smaller eyes.

**Fig. 3.**
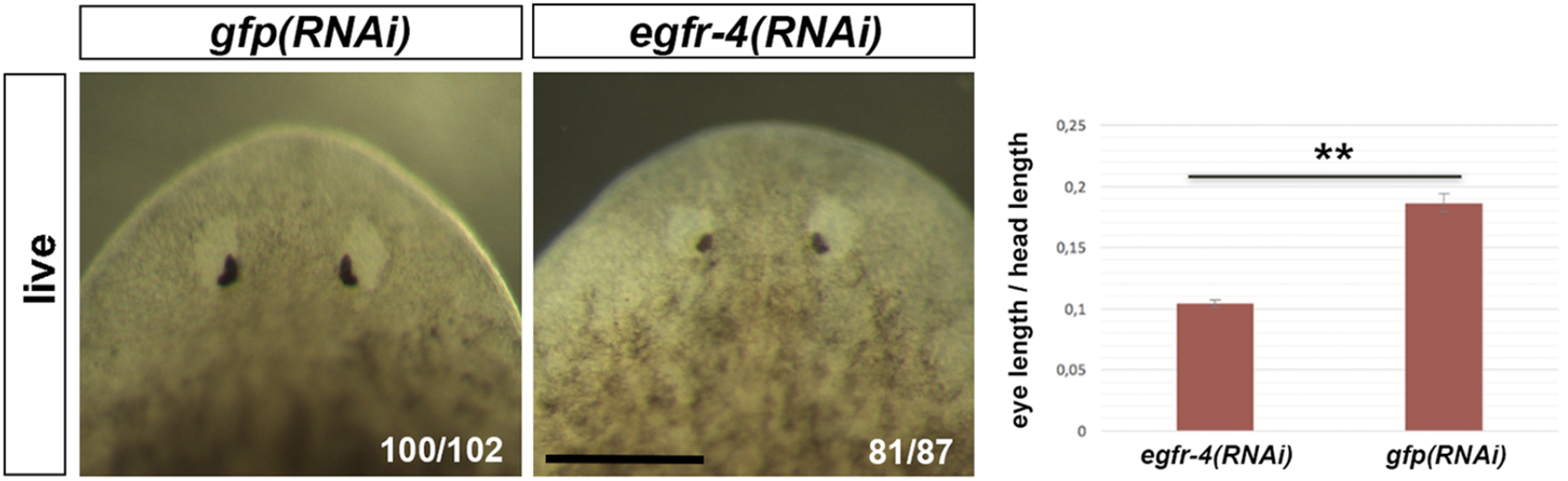
*Smed-egfr-4(RNAi)* planarians regenerate smaller eyes. The silencing of *Smed-egfr-4* results in smaller eyes during anterior regeneration. The graph shows the result of quantifying the ratio between the eye length and the length of the head region from the base of the eye to the head anterior tip. n=40 eyes measured for each treatment. **p<,005 (t-test). All samples correspond to 11 days of regeneration. Scale bar, 200 μm.

### Smed-egfr-4(RNAi) animals regenerated lower numbers of mature eye cells but a higher number of eye progenitors

As externally *Smed-egfr-4(RNAi)* planarians regenerated smaller eyes we analysed whether this reduction corresponded to a lower number of eye cells. VC-1 is an anti-arrestin antibody that recognizes specifically planarian photoreceptor neurons (Sakai et al., 2000). On the other side, *Smed-tph* codes for a tryptophan hydroxylase homologue that is expressed in the pigmented eye-cup cells (Fraguas et al., 2011). The cell bodies of the photoreceptor neurons are all clustered around the pigmented eye-cup. In order to quantify the number of photoreceptors and eye-cup cells we performed double *in situ* hybridizations and immunostainings for *Smed-tph* and VC1, respectively (Fig. 4 A-H). Cell quantifications showed that both cell types were significantly reduced after *Smed-egfr-4* RNAi (Fig. 4 I-J). It has been reported that *Smed-ovo* labels both mature eye cells as well as eye progenitor cells (Lapan and Reddien, 2012). Eye progenitor cells are localized as two trails of cells posterior to the mature eyes. It has been described that those progenitors differentiate as they approach the mature eye (Lapan and Reddien, 2012). In order to determine whether the reduction in the number of mature eye cells observed after the silencing of *Smed-egfr-4* was caused by a reduction in the number of eye progenitors we analysed the expression of *Smed-ovo* after *Smed-egfr-4* RNAi. Remarkably, and in contrast to what happens for the mature cells, we observed an increase in the number of *Smed-ovo+* cells in those animals (Fig. 4 K-L). In controls *Smed-ovo+* cells were mainly located between the mature eyes and the base of the pharynx. However, after the silencing of *Smed-egfr-4, Smed-ovo+* cells were observed in more posterior regions (Fig. 4 K-L). We quantified the number of *Smed-ovo+* cells and found a significant increase of these cells in *Smed-egfr-4(RNAi)* animals (Fig. 4M). Whereas *Smed-ovo* is expressed in progenitors of both photoreceptor neurons and pigment eye-cup cells, *Smed-sp6-9* is expressed in the progenitors of the pigment eye-cup cells (Lapan and Reddien 2011, 2012). After the silencing of *Smed-egfr-4,* the number of *Smed-sp6-9* cells also increased significantly (Fig. 4 N-P). These results indicate that the specification of pluripotent neoblasts into eye progenitors is not affected after *Smed-egfr-4* RNAi but suggest that the final differentiation of these progenitors into mature eye cells is compromised.

**Fig. 4.**
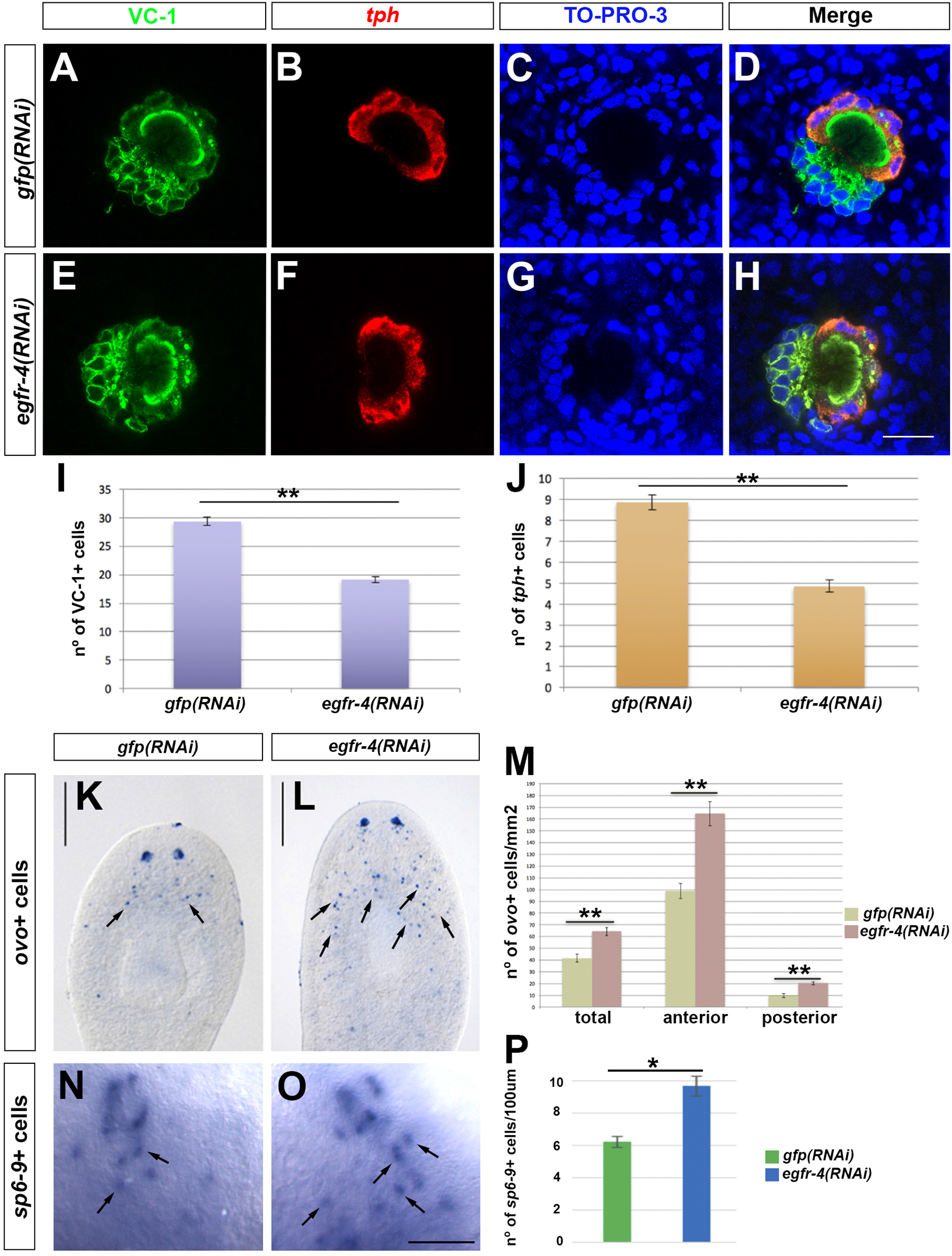
Reduced numbers of mature eye cells and higher number of eye progenitors after *Smed-egfr-4 RNAi.* (A-H) Double fluorescent *in situ* hybridization for *Smed-tph* and immunostaining with VC-1 to detect the pigmented eye-cup (in red) and the photoreceptor neurons (in green) in control (A-D) and Smed-egfr-4(RNAi) animals (EH). Nuclear staining in blue with TO-PRO-3. Scale bar, 20 μm. (I, J) Graphs showing the quantification of VC-1+ and tph+ cells, respectively. In (I) n=23 eyes for each treatment. In (J) n=15 eyes for each treatment **p<0,005 (t-test). All samples correspond to 11 days of regeneration. (K,L) Whole mount *in situ* hybridization for *Smed-ovo.* Arrows point to eye progenitors in the pre-pharyngeal region. Scale bar, 300 μm. (M) Graph of the quantification of ovo+ cells all throughout the animal. Total refers to the whole animal; anterior refers to the region from the tip of the head to the base of the pharynx; posterior refers to the length between the base of the pharynx and the posterior tip of the tail. All samples correspond to 10 days of regeneration. n=20 animals for each treatment. **p<0,005 (t-test). (N,O) Whole-mount *in situ* hybridization for *Smed-sp6-9.* Arrows point to eye progenitors in the pre-pharyngeal region. Scale bar, 40 μm. (P) Graph of the quantification of *sp6-9+* cells. All samples correspond to 11 days of regeneration. n= 25 animals for each treatment. *p<0,05 (t-test).

## Discussion

Recent advances are turning planarians as an excellent model in which to study stem cell differentiation *in vivo.* Planarian pluripotent stem cells specialize into lineage-committed progenitors that will further differentiate into all the cell types found in these animals (Scimone et al., 2014; Zhu and Pearson, 2016). Not much, however, is known yet about how this differentiation process is regulated at the molecular level. Still, few genes have been reported to have a function during neoblast differentiation. Thus, for example, *Smed-mex-3* is required for the specification of the different progenitor populations (Zhu et al., 2015); also, the silencing of *Smed-not* impairs neoblast differentiation (Solana et al., 2013). Planarian eyes are constituted by two cell types: pigmented cells that organize themselves into a pigmented eye-cup and bipolar photoreceptor neurons than send rhabdomeric dendrites to the eye-cup and visual axons to a specific region of the brain. Moreover, some visual axons project contralaterally crossing the midline and forming an optic chiasm (Sasaki et al., 2000; Okamoto et al., 2005). Recently, a transcriptomic analysis of planarian eyes has characterized several genes involved in the differentiation process from pluripotent neoblast to eye progenitors and then to mature eye cells. The earliest progenitors committed towards the eye lineage co-express *Smed-ovo, Smed-six-1/2* and *Smed-eya.* The silencing of any of these genes inhibits eye regeneration (Pineda et al., 2000; Mannini et al. 2004; Lapan and Reddien 2012). From these *ovo+* early progenitors it appears that both lineages, photoreceptor neurons and pigmented eye-cup cells, split (Lapan and Reddien, 2012). Also, among photoreceptor neurons not all of them are equivalent as several subpopulations (anterior, dorsal posterior and ventral posterior) can be distinguished by the expression of specific markers (Collins et al., 2010). Interestingly, it has been described that *Smed-smad6/7-1* and *Smed-bmp* are involved in the differentiation of the anterior photoreceptor neurons (González-Sastre et al., 2012).

Here, we have addressed the role of *Smed-egfr-4* in planarian regeneration. Despite being expressed in the CNS, the silencing of this gene does not result in any defect in its regeneration, which might be indicating that other EGFRs could be compensating for the lack of *Smed-egfr-4* function. Alternatively, the silencing of *Smed-egfr-4* could result in defects in the CNS that cannot be detected with the actual available markers. On the other hand, the silencing of *Smed-egfr-4* results in a significant decrease in the number of mature eye cells of both types. Remarkably, this decrease is coupled with an increase in the number of *Smed-ovo+* and *Smed-sp6-9+* progenitor cells located at the pre-pharyngeal region. In a current model for eye differentiation, pluripotent neoblasts (cNeoblasts) give rise to early eye progenitors defined by the co-expression of *ovo, eya* and *six-1/2* (Lapan and Reddien, 2012). From them the two eye lineages split and generate late eye progenitors. Those expressing *ovo, sp6-9* and other factors give rise to the pigmented eye-cup cells whereas those expressing *ovo* and other factors give rise to the photoreceptor neurons (Lapan and Reddien, 2012). The fact that the final differentiation of both lineages, photoreceptor neurons and pigmented eye-cup cells, are affected and that we observe an increase of *ovo+* and *sp6-9+* cells suggest that *Smed-egfr-4* is probably required for the differentiation of both lineages once they have diverged from a common early progenitor (Fig. 5). These results on the function of *Smed-egfr-4* on eye progenitor differentiation resemble those on the role of *Smed-egfr-1* on the differentiation of the gut progenitors (Barberán et al., 2016a) and support the hypothesis that in planarians the EGFR signalling pathway could play a general role on the differentiation of lineage-committed progenitors (Barberán and Cebrià, 2018). Further studies should try to characterize additional factors involved in the final differentiation of late eye progenitors as well as the role of other EGFRs in the differentiation of other planarian cell types.

**Fig. 5.**
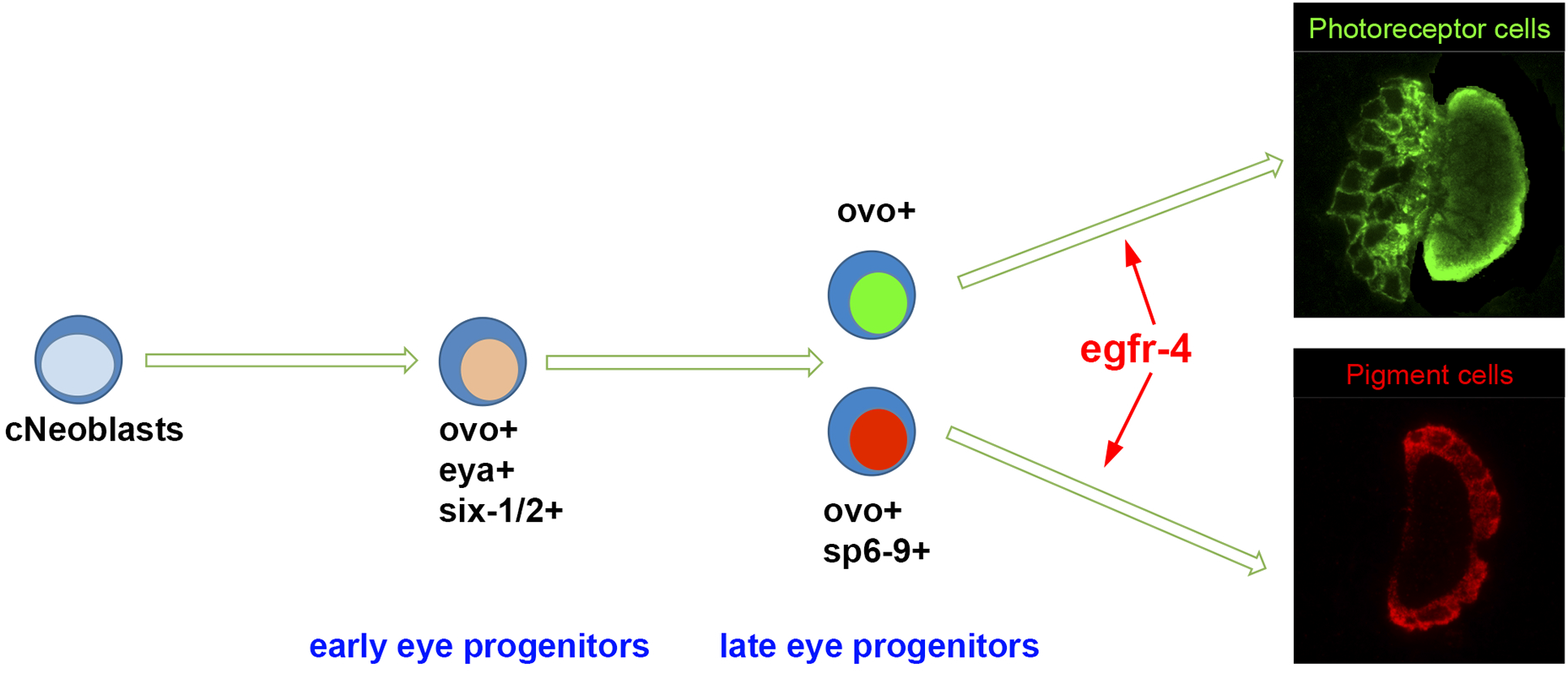
Proposed model for the role of *Smed-egfr-4* during eye regeneration. See details within the main text.

## Material & Methods

### Planarian culture

Asexual *Schmidtea mediterranea* from the BCN-10 clonal line was used in these experiments. Animals were fed with veal liver and starved for at least 1 week before all experiments.

### RNA interference

Silencing by RNAi (RNA interference) was performed as previously described (Sánchez Alvarado and Newmark, 1999). Control animals were injected with double-stranded RNA (dsRNA) of green fluorescent protein (GFP). All animals went through two rounds of dsRNA injection and amputation, separated by 3-4 days.

### *In situ* hybridization and immunohistochemistry

For whole mount *in situ* hybridizations animals were treated as previously described (King and Newmark, 2013). Fluorescent *in situ* hybridization for *Smed-tph* was followed by immunostaining with the monoclonal antibody VC-1 against photoreceptor neurons (Sakai et al., 2000), diluted 1:15,000. A goat anti-mouse secondary antibody conjugated to Alexa 488 (Molecular Probes) was used at a 1:400 dilution. Nuclear staining was achieved by using TO-PRO-3. Non-fluorescent in situ hybridizations were imaged with a Leica MZ16F stereomicroscope and a ProgRes C3 camera (Jenoptik). Fluorescence images were obtained with a Leica SPE confocal microscope. All images were analysed and treated with ImageJ and Photoshop.

### Statistical analyses

Statistical analyses were performed using MS Excel and R Software 3.5-0 (RStudio 1.1.453). All the results for the graphs are expressed as the mean ± standard error of the mean (s.e.m.). Data were analysed with t-test and a p-value below 0,05 was considered statistically significant. All the quantifications were performed by a blind-observer.

## Acknowledgements

We would like to thank all members of the lab, especially Susanna Fraguas and Sara Barberán for their help and technical support. This work was funded by grant BFU2015-65704P from Ministerio de Economía y Competitividad (Spain) to F.C.

## Abbreviations

*EGFR*: epidermal growth factor receptor
*CNS*: central nervous system RNAi
*RNAi*: RNA interference

## References

Barberán, S., Fraguas, S., Cebrià, F. (2016a). The EGFR signaling pathway controls gut progenitor differentiation during planarian regeneration and homeostasis.Development 143: 2089–2102.

Barberán, S., Martín-Durán, J.M., Cebrià, F. (2016b). Evolution of the EGFR pathway in Metazoa and its diversification in the planarian Schmidtea mediterranea.Sci Rep 6: 28071.

Barberán, S., Cebrià, F. (2018). The role of the EGFR signalling pathway in stem cell differentiation during planarian regeneration and homeostasis.Semin Cell Dev Biol pii: S1084–9521(18)30032-6. doi: 10.1016/j.semcdb.2018.05.011.

Cebrià, F., Adell, T., SALÓ, E. (2018). Rebuilding a planarian: from early signalling to final shape. Int J Dev Biol 62: 537–550.

Collins, J.J. 3rd, Hou, X., Romanova, E.V., Lambrus, B.G., Miller, C.M., Saberi, A., Sweedler, J.V., Newmark, P.A. (2010). Genome-wide analyses reveal a role for peptide hormones in planarian germline development. PLoS Biol 8: e1000509.

Fraguas, S., Barberán, S., Cebrià, F. (2011). EGFR signaling regulates cell proliferation, differentiation and morphogenesis during planarian regeneration and homeostasis. Dev Biol 354: 87–101.

González-Sastre, A., Molina, M.D., SalÓ, E. (2012). Inhibitory Smads and bone morphogenetic protein (BMP) modulate anterior photoreceptor cell number during planarian eye regeneration. Int J Dev Biol 56: 155–163.

King, R.S., Newmark, P.A. (2013). In situ hybridization protocol for enhanced detection of gene expression in the planarian Schmidtea mediterranea. BMC Dev Biol 13:8.

Lapan, S.W., Reddien, P.W. (2011). dlx and sp6-9 control optic cup regeneration in a prototypic eye. PLoS Genet 7: e1002226.

Lapan, S.W., Reddien, P.W. (2012). Transcriptome analysis of the planarian eye identifies ovo as a specific regulator of eye regeneration. Cell Rep 2: 294–307.

Mannini, L., Rossi, L., Deri, P., Gremigni, V., Salvetti, A., Saló, E., Batistoni, R. (2004). Djeyes absent (Djeya) controls prototypic planarian eye regeneration by cooperating with the transcription factor Djsix-1. Dev Biol 269: 346–359.

Okamoto, K., Takeuchi, K., Agata, K. (2005). Neural projections in planarian brain revealed by fluorescent dye tracing. Zoolog Sci 22: 535–546.

Pineda, D., González, J., Callaerts, P., Ikeo, K., Gehring, W.J., Saló, E. (2000). Searching for the prototypic eye genetic network: Sine oculis is essential for eye regeneration in planarians. Proc Natl Acad Sci USA 97: 4525–4529.

Rink, J.C. (2018). Stem cells, patterning and regeneration in planarians: selforganization at the organismal scale. Methods Mol Biol 1774: 57–172.

Rink, J.C., Vu, H.T., SáNchez-Alvarado, A. (2011). The maintenance and regeneration of the planarian excretory system are regulated by EGFR signaling. Development 138: 3769–3780.

Sakai, F., Agata, K., Orii, H., Watanabe, K. (2000). Organization and regeneration ability of spontaneous supernumerary eyes in planarians —eye regeneration field and pathway selection by optic nerves-. Zoolog Sci 17: 375–381.

SáNchez-Alvarado, A., Newmark, P.A. (1999). Double-stranded RNA specifically disrupts gene expression during planarian regeneration. Proc Natl Acad Sci USA 96: 5049–5054.

Scimone, M.L., Kravarik, K.M., Lapan. S.W., Reddien, P.W. (2014a). Neoblasts specialization in regeneration of the planarian Schmidtea mediterranea. Stem Cell Reports 3, 339–52.

Solana, J., Gamberi, C., Mihaylova, Y., Grosswendt, S., Chen, C., Lasko, P., Rajewsky, N., Aboobaker, A. (2013). The CCR4-NOT complex mediates deadenylation and degradation of stem cell mRNAs and promotes planarian stem cell differentiation. PLoS Genet 9: e1004003.

Tu, K.C., Cheng, L.C., Vu, H.T., Lange, J.J., McKinney, S.A., Seidel., C.W., Sánchez-Alvarado, A. (2015). Egr-5 is a post-mitotic regulator of planarian epidermal differentiation. eLife 4: e10501.

Wagner, D.E., Wang, I.E., Reddien, P.W. (2011). Clonogenic neoblasts are pluripotent adult stem cells that underlie planarian regeneration. Science 332: 811–6.

Zeng, A., Li, H., Guo, L., Gao, X., McKinney, S., Wang, Y., Yu, Z., Park, J., Semerad, C., Ross, E., Cheng, L.C., Davies, E., Lei, K., Wang, W., Perera, A., Hall, K., Peak, A., Box, A., SáNchez Alvarado, A. (2018). Prospectively isolated tetraspanin+ neoblasts are adult pluripotent stem cells underlying planaria regeneration. Cell 173: 1593–1608.

Zhu, S.J., Pearson, B.J. (2016). (Neo)blast from the past: new insights into planarian stem cell lineages. Curr Opin Genet Dev 40: 74–80.

Zhu, S.J., Hallows, S.E., Currie, K.W., Xu, C., Pearson, B.J. (2015). A mex3 homolog is required for differentiation during planarian stem cell lineage development. eLife 4: e07025.

